# The permeation of potassium ions through the lipid scrambling path of the membrane protein nhTMEM16

**DOI:** 10.1101/2022.03.21.485163

**Authors:** Xiaolu Cheng, George Khelashvili, Harel Weinstein

## Abstract

The TMEM16 family of transmembrane proteins includes Ca^2+^-activated phospholipid scramblases (PLS) that can also function as non-selective ion channels. Extensive structural and functional studies have established that a membrane-exposed hydrophilic groove in TMEM16 PLS can serve as a translocation pathway for lipids. However, it is still unclear how the TMEM16 PLS conduct ions. A “protein-delimited pore” model suggests that ions are translocated through a narrow opening of the groove region, which is not sufficiently wide to allow lipid movement, whereas a “proteolipid pore” model envisions ions and lipids translocating through an open conformation of the groove. We investigated the dynamic path of potassium ion (K^+^) translocation that occurs when an open groove state of nhTMEM16 is obtained from long atomistic molecular dynamics (MD) simulations and calculated the free energy profile of the ion movement through the groove with umbrella sampling methodology. The free energy profile identifies effects of specific interactions along the K^+^ permeation path. The same calculations were performed to investigate ion permeation through a groove closed to lipid permeation in the nhTMEM16 L302A mutant which exhibits a stable conformation of the groove that does not permit lipid scrambling. Our results identify structural and energy parameters that enable K^+^ permeation and suggest that the presence of lipids in the nhTMEM16 groove observed in the simulations during scrambling or in/out diffusion, affect the efficiency of K^+^ permeation to various extents.

## Introduction

The TMEM16 (Anoctamin) family proteins are involved in a wide range of physiological processes in eukaryotic organisms. Some family members, such as TMEM16A ^1^ and TMEM16B ^2^, are Ca^2+^-activated Cl^−^ channels required for normal electrolyte and fluid secretion in various key physiological functions, e.g., mediating olfactory amplification ^2^. Others, such as TMEM16F and TMEM16K, have dual functions as Ca^2+^-activated phospholipid scramblases and non-selective ion channels ^3-5^. While not all the physiological functions of mammalian TMEM16 proteins have been elucidated thus far, mutations have been associated with various disorders ^6,7^. Inferences from the structure of various TMEM16 family members determined with X-ray or cryo-electron microscopy (cryo-EM) ^5,8-16^ have reached important mechanistic insights into both ion conduction and lipid scrambling functions by these proteins. A polar groove identified in the X-ray structure of the two protomers of the fungal scramblase nhTMEM16 ^8^ was first proposed to be the lipid translocation pathway (Figure 1). Subsequent structural studies of nhTMEM16 constructs showed that the groove adopts different conformations in the presence and absence of Ca^2+^, including an intermediate conformation between the open and closed forms ^12,14^. The structure of TMEM16A, a mammalian Ca^2+^-activated Cl^−^ channel, revealed a closed groove in the presence of Ca^2+ 9-11^. In contrast, afTMEM16 scramblase, another fungal homologue, was found to adopt an open-groove conformation in the presence of Ca^2+^ even in a membrane environment in which the scrambling activity is low ^13^. Of the mammalian scramblases, the structure of mTMEM16F ^15,16^ has thus far been determined only with a closed groove in the presence of Ca^2+^, while TMEM16K, a scramblase resident in the endoplasmic reticulum, exhibits open and closed conformations of the groove in the presence and absence of Ca^2+^, respectively ^5^.

**Figure 1:**
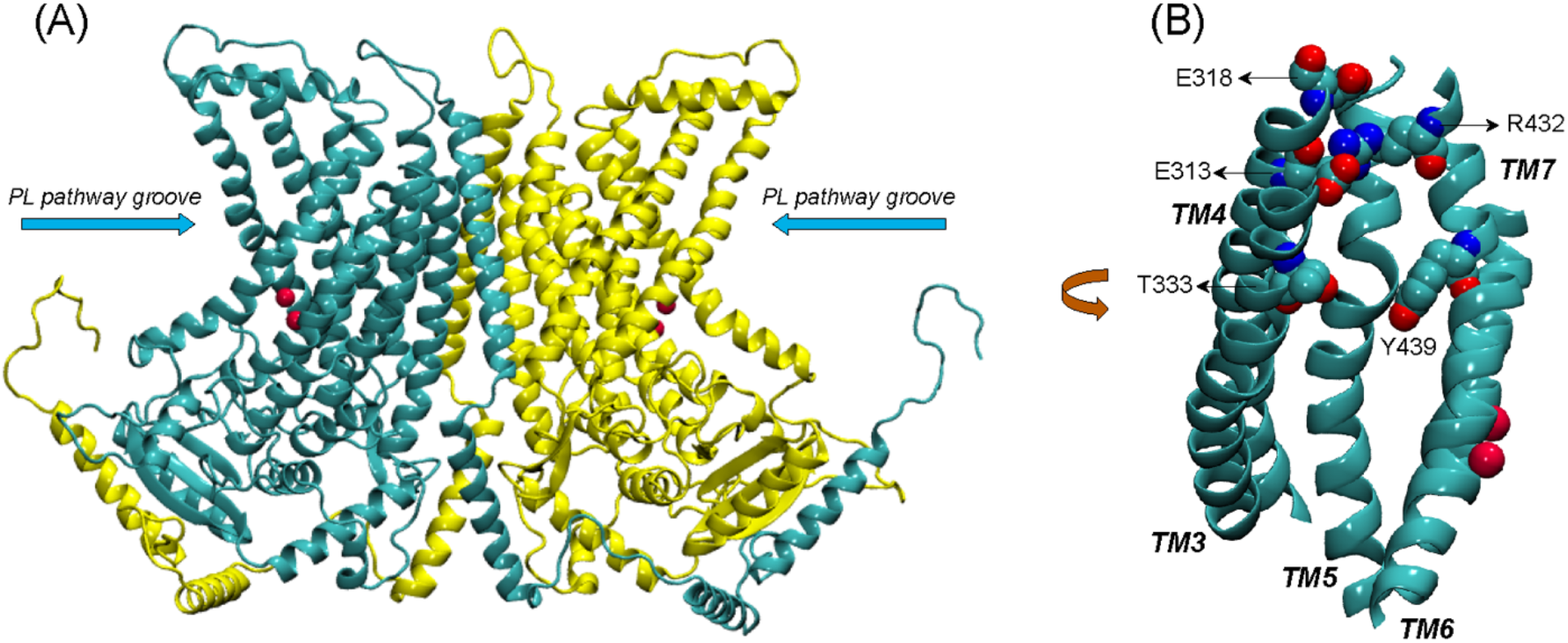
(**A**) Complete structural model of nhTMEM16 dimer (based on PDBID 4WIS). The two monomers of the dimer are indicated in cyan and yellow color, respectively. The phospholipid (PL) pathway groove region is indicated, and the bound Ca^2+^ ions (two per each monomer) are shown as red spheres. (**B**) View of the groove area (TMs 3-7) of one of the monomers, highlighting the EC network of polar residues (E313, E318, R432, T333, and Y439).

Computational studies using molecular dynamics (MD) simulations based on the structural information available for the various TMEM16 family members ^5,14,17-20^ have aimed to complete the mechanistic picture of lipid and ion translocations by these proteins. Results from our previous µs-long all-atom MD studies of structure-function relations of nhTMEM16 ^14,19,20^ have shown that the network of polar residues on the extracellular side of the groove region (residues R432/E313/E318 and Y439/T333, see Figure 1) functions as gates that modulate the exchange of lipids between the groove and the bulk membrane. The long simulations trajectories also revealed events of ion passage through the groove in the absence of artificially applied voltage across the membrane. Here we pursue this intriguing opportunity to investigate the molecular mechanisms associated with ion permeation in these scramblases in the structural context of the conformations visited in the dynamics of the nhTMEM16 scramblase.

Early data have led to the suggestion that ion channel activity of the nhTMEM16 scramblase is mediated by a protein-delimited pore generated according to a proposed “alternating pore-cavity” model ^12,14^. Thus, structure determination showed that a much narrower opening of the groove region in an intermediate conformation can create a protein-enclosed pore large enough to permit the permeation of ions but not lipids. Indeed, we found that the single mutation L302A of nhTMEM16 stabilizes such an intermediate conformation, and the assays of lipid scrambling and ion flux by this mutant supported the “alternating pore-cavity” model ^14^. On the other hand, results from modeling and computational simulations of electrophysiology assays of this scramblase ^21^ support a proteolipidic-pore model, which assumes that ions and lipids are translocated through a single protein conformation. We report here the investigation of the dynamic path of potassium ion (K^+^) translocation through the open groove of nhTMEM16 observed in the long MD simulations, and present the free energy profile of the ion movement through the groove calculated with umbrella sampling methodology ^22^. To investigate ion permeation in a model with a groove closed to lipid permeation, the same calculations are performed with the MD simulation trajectories of the nhTMEM16 L302A mutant which exhibits a stable conformation of the groove that does not permit lipid scrambling ^14^. Our results quantify the energetic feasibility of K^+^ permeation through different conformations of the groove and the slowing down of the process when membrane lipids occupy the groove.

## Results

### The translocation pathway of K^+^ through WT nhTMEM16 includes specific interaction sites in the groove

We have previously reported unbiased 10 *µ*s all-atom MD simulations of nhTMEM16 (PDBID 4WIS ^8^) in 3:1 POPE/POPG membrane system immersed in a solution containing 150 mM K^+^Cl^- 14^ (see Methods for details). The trajectory from this MD simulation contains two separate events of ion translocating through the protein groove recorded in Supplementary files: movie 1 and movie 2. In each case, the translocated ion was a potassium ion (K^+^). Snapshots representing various stages of the ion translocation during the two events are shown in Figure 2.

**Figure 2:**
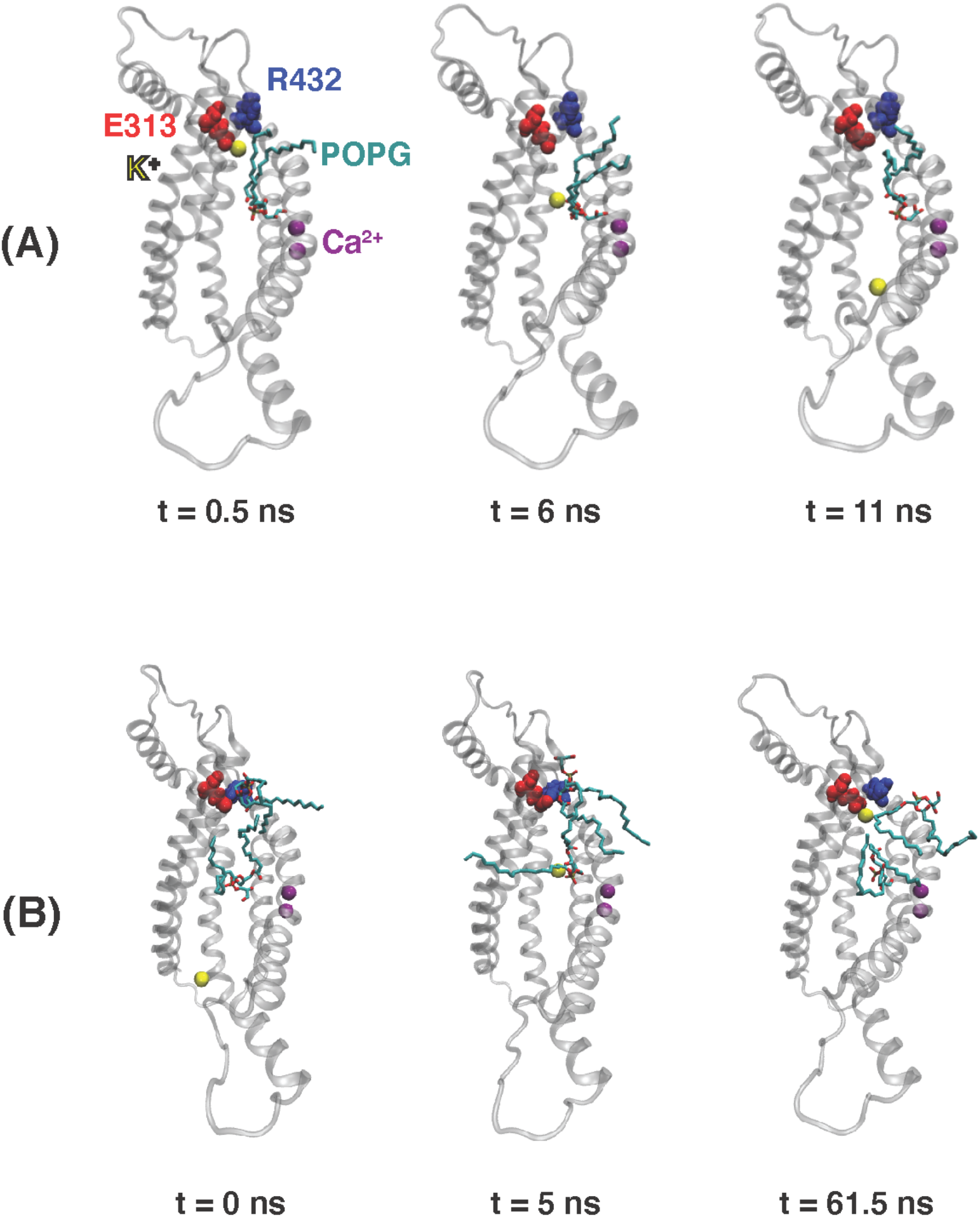
Snapshots from the 10 µs MD simulation of nhTMEM16 capturing the potassium ion permeation events in Supplementary movies 1 and 2. Time zero in the figure corresponds to the beginning of each described translocation processes occurring at 5.8 µs and 7 µs, respectively (see also Fig 3A-B below) **(A)** Permeation from EC to IC. **(B)** Permeation from IC to EC. Note that for the EC-to-IC translocation process (panel A) we show the first frame at t = 0.5 ns, as the ion is distant from the groove at t = 0 which is the starting frame for the Supplementary movie 1. The potassium ion being translocated is shown as yellow sphere, POPG lipids are shown as licorice, TM3–TM7 of nhTMEM16 are rendered in transparent cartoon, E313 is shown as red spheres, R432 is shown as blue spheres, and Ca2+ ions are shown as purple spheres.

The K^+^ ion permeation occurring at the 5.8 µs time-point in the MD trajectory shown in Supplementary movie 1 is from the extracellular (EC) to the intracellular (IC) end. In the other recorded translocation event – at 7 µs in the trajectory (Supplementary movie 2), a K^+^ traverses the groove in the IC-to-EC direction. The duration of the permeation events was 11 ns for the first and 63 ns for the second one (see panels A and B in Figure 3). As can be seen from these snapshots and the Supplementary movies, the translocated K^+^ ion forms interactions at two main sites in the groove during both events. These are i) the EC gate ^19^ formed by the E313-R432 residue pair, and ii) a POPG lipid situated in the middle of the groove and coordinated by several residues (Supplementary Figure S1). Specifically, during the EC-to-IC permeation event, the EC gate is broken (Figure 3G) and the incoming K^+^ ion partitions at E313 (Figure 3C), which is disengaged from R432. After spending 3 ns at E313 (Figure 3C), the K^+^ ion continues down the groove and engages in electrostatic interactions with the headgroup of a POPG lipid (Figure 3E) that was seen to penetrate the groove from the IC side at earlier time-point in the trajectory (Supplementary Figure S2). From this interaction site the ion continues to travel down the groove and diffuses away into the solution.

**Figure 3:**
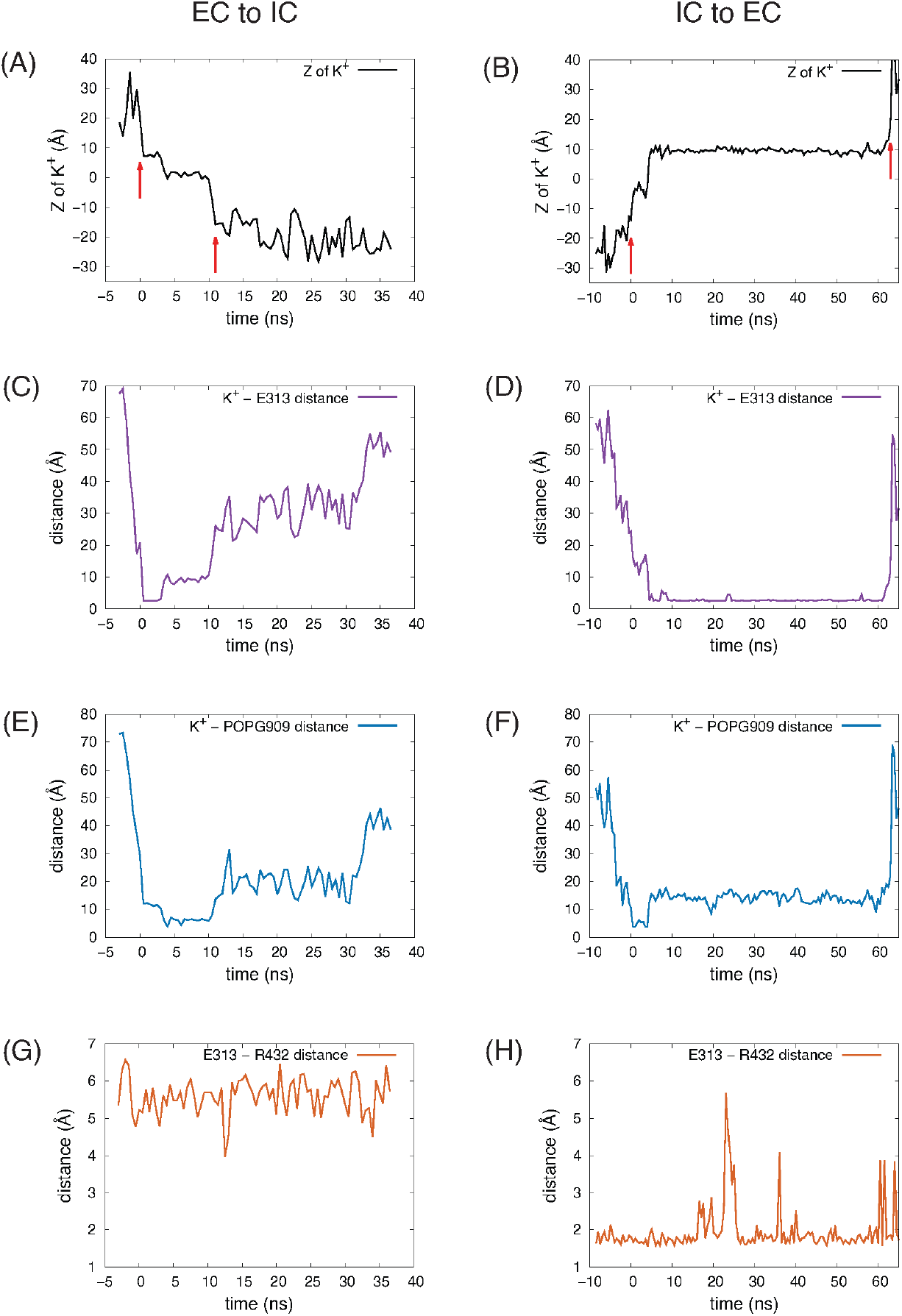
Dynamic changes during the K+ ion translocation events in the 10µs unbiased MD simulation of nhTMEM16: (**A-B**) Changes in the Z position of the translocated K^+^ ion during the EC-to-IC (**A**) and IC-to-EC (**B**) permeation events. Red arrows indicate the beginning and end of the translocation process. The beginning of the EC-to-IC process (t=0 was defined by the time-point in the trajectory when the ion partitioned from the bulk into the groove (established interaction with E313), while the end of the IC-to-EC process was defined by the time-point when the ion disengaged from E313 and diffused away to the EC milieu (See Supplementary movies). Z =0 Å represents the position of the C_α_ atom of T381. (**C-D**) Distance between the translocated K^+^ ion and E313. (**E-F**) Distance between the translocated K^+^ ion and the closest headgroup P atom of the in-groove POPG lipid. (**G-H**) Distance between the EC gate residues, E313 and R432. The dynamics are presented in the Supplementary movies which start at the t = 0 ns time point.

As observed also for the scrambling events recorded in our MD simulations ^19^, the K^+^ ion translocation pathway in the IC-to-EC direction mirrors the one described above, but in reverse order. Specifically, the K^+^ ion engages first with the same POPG lipid in the groove as described above for the EC-to-IC event (Figure 3F), and then proceeds towards the EC gate which opens transiently as K^+^ remains engaged with E313 (Figure 3D and 3H). After interacting with E313 for 58 ns (Figure 3D), the ion diffuses into the EC milieu completing the translocation (Figure 3B). During this time, a network of interactions was observed among four sites: the potassium ion, E313, R432 and another POPG in the EC region.

Together, the simulation results suggest a mechanistic role for the electrostatic interactions between K^+^ ions and the two sites along the groove (the E313 residue, and the groove-bound POPG lipid), akin to the ion-binding sites in ion channels that are considered to serve as “stepping stones” for the ions to pass through the conducting pore ^17,23^. The involvement of the groove-bound POPG lipid in the permeation sequence is particularly intriguing, since it has been debated whether lipids and ions share the same translocation pathway in the TMEM16 scramblases and thus can be translocated concomitantly, or the lipid scrambling and ion permeation events use different pathways and thus are mechanistically decoupled ^12,14^. We find that the POPG lipid that associates with the translocated K^+^ ion was not scrambled itself (Supplementary Figure S2), but its strategic location in the middle of the groove region and its involvement in the ion translocation in both directions, suggest that the negative charge of its headgroup may lower the free energy barrier for the K^+^ ion permeation process. To test this hypothesis, we used umbrella sampling simulations ^22^ to calculate the free energy of K^+^ ion translocation in the presence or absence of the groove-bound POPG lipid, as well as under conditions when the negatively charged PG headgroup of the bound lipid was substituted by a zwitterionic PC headgroup. The umbrella sampling simulation of wild type nhTMEM16 were done under four different conditions (see Methods): System 1 – the original system in which the two K^+^ permeation events were observed, consisting of the scramblase with the POPG lipid in the groove, embedded in a 3:1 POPE/POPG membrane; System 2 – the same a System 1 but without the in-groove POPG; System 3 – a system consisting of the scramblase with a POPC in the groove (instead of POPG), embedded in a POPC membrane; System 4 – the same a System 3 but without the in-groove POPC. The headgroup of the in-groove POPC in System 3 is at the same Z position along the membrane axis as the head group of the in-groove POPG of System 1 (see Supplementary Figure S3). System 2 and System 4 are without lipid in the groove, but they differ in the membrane composition as detailed above.

### The potential of mean force profile for the translocation of K^+^ in the open groove of nhTMEM16

The PMF in each of the four Systems was calculated from the umbrella sampling simulations along the reaction coordinate taken as the distance in the Z direction (perpendicular to the membrane) between the K^+^ ion and the C_α_ atom of residue T381 (see Methods). The resulting free energy curves are shown in Figure 4. The most striking feature is the large difference between the energy barrier calculated for System 3 compared to those of Systems 1, 2, and 4. While the maxima occur at very similar Z values in all Systems, the one in System 3 is ∼7 kcal/mol while in all the others are ∼2 kcal/mol. This suggests that the in-groove presence of the large headgroup of the POPC lipid molecule hinders the translocation, likely by steric obstruction. Notably, the curve representing results for System 1 has a local minimum near the groove center, at Z = ∼2.7 Å where the curve for System 3 has its maximum. This reflects the favorable electrostatic interaction of the permeating K^+^ ion with the negatively charged headgroup of POPG.

**Figure 4:**
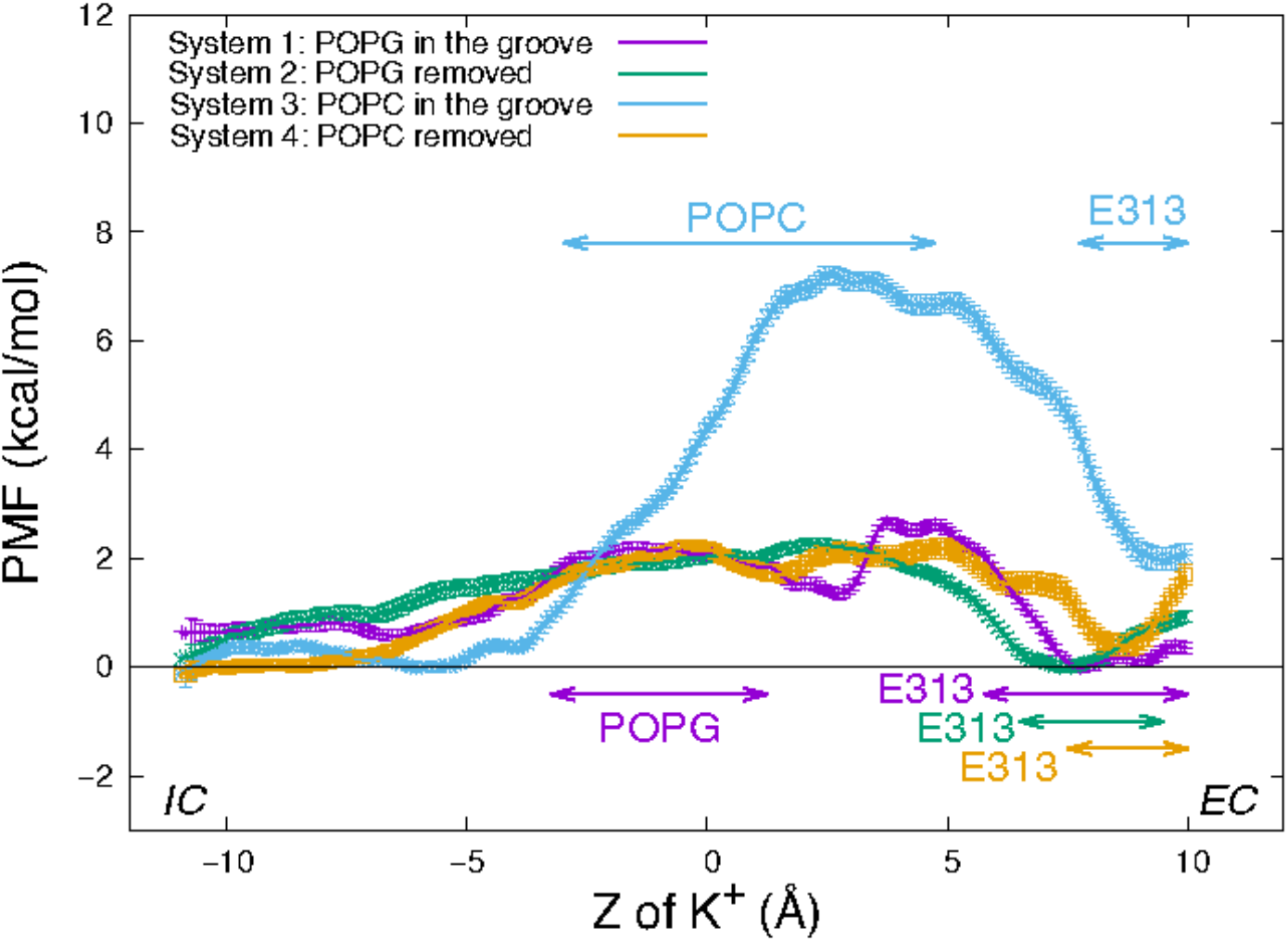
Potential of mean force of a K+ ion along the permeation pathway in the four wild type nhTMEM16 Systems (see text). The PMF profiles obtained for each System are depicted with error bars calculated as described in Methods. Z = 0 is set to be the Z position the Cα atom of T381 in the protomer. The ranges of “critical interactions” in which the K^+^ ion is engaged in >50% of the trajectory frames in a PMF window are marked by horizontal double-arrowed lines with System-corresponding colors.

However, the similarity between the free energy barriers for Systems 1 and 2 suggests that the free energy contribution from the favorable electrostatic attraction between the ion and POPG lipid at a longer distance is compensated at shorter interaction distances by unfavorable steric interactions due to the presence of the POPG lipid in the groove. The greater similarity between the PMF profiles of System 2 and 4 is expected as the translocating K+ ion samples similar (open) groove conformations without lipid present in the middle of the groove.

The nature of the interactions shaping the PMF profiles in Figure 4 is brought to light by relating it to the number of trajectory frames in which a residue or lipid is within 3Å of the translocated ion. Results of this analysis performed separately for each umbrella sampling window from the equilibrated umbrella sampling simulation are illustrated in Table 1 in which the results are shown for the System 1 calculations in the umbrella window centered near the POPG lipid located in the groove. Based on such data obtained for each umbrella window we identified as “critical interactions” those in which the K+ ion is engaged in >50% of the trajectory frames in that window. Thus, for the umbrella window shown in Table 1, the POPG lipid emerges as such a critical interaction partner. The PMF energy profiles in **Figure 4** identify effects of other specific interactions along the K^+^ permeation path, assigned to regions of the PMF profiles in Figure 4. These assignments are indicated for each of the four Systems by the color code of the horizontal double arrows. The different lengths of the double arrows correspond to the time lengths of the interactions (see Supplementary Figures S4-S7 for a complete annotation of the PMF profiles). In all four Systems, the interval along the reaction coordinate where the K+ ion interacts with E313 coincides with the local minimum at the EC end in the PMF profile, suggesting that E313 serves as an attraction point for the ion at the EC end of the groove. The PMF maximum (7 kcal/mol peak) for the in-groove POPC construct, System 3, is located in the interval where the K+ ion interacts with the POPC lipid (from Z = -3 Å to Z = 4.75 Å), suggesting that indeed the presence of POPC in the groove results in the free energy barrier that is ∼5 kcal/mol higher than in the other three systems.

**Table 1:** Illustration of frame-counting results for the 3Å proximity to K^+^ in an umbrella sampling window (the center of the window Z_0_ is -1.5 Å away in Z direction from the C_α_ atom of residue T381) in System 1. The total number of frames in the PMF windows is 4500 and an interaction with the potassium ion and is considered “critical” if it is present in >2250 frames (i.e., > 50%). Note, however that the total number of interactions is >4500 because in a single frame, the K+ ion can interact with several residues. Thus, the sum of percentage in the second column is not 100%.

| $Z_0 = -1.5 \text{ \AA}$ | ~~4500~~ total frames 4939 |
| --- | --- |
| Residue | Number of “interacting” frames |
| POPG 909 | 3732 (82.9%) |
| THR 381 | 953 (21.2%) |
| VAL 337 | 135 (3.0%) |
| SER 382 | 53 (1.2%) |
| TYR 513 | 37 (0.8%) |
| THR 340 | 18 (0.4%) |
| LEU 336 | 9 (0.2%) |
| PHE 440 | 2 (0.04%) |

Interestingly, the interval of the K^+^/POPC interactions in System 3 is longer than for the K^+^/POPG interactions (from Z = -3.25 Å to Z = 1.25 Å) in System 1. This difference could be due to the smaller size of the PG head group compared to PC. Notably, this short K^+^/POPG interacting range does not include the local minimum at Z = 2.7 Å in the PMF profile for System 1. Rather, this minimum appears to be due to the combined longer range electrostatic interaction as well as a favorable environment generated by the polar residues T381 and S382 (see Supplementary Figure S4) located just above the POPG lipid binding site (Supplementary Figure S8).

### A narrowing of the groove leads to higher free energy barriers for K^+^ ion translocation in the L302A mutant of nhTMEM16

We recently described the L302A mutant of nhTMEM16 in which the groove is significantly narrower, which abrogates lipid scrambling while retaining ion channel activity ^14^. The much narrower opening of the groove region is seen to generate a protein-enclosed pore that can sustain the translocation of certain ions but not lipids. To determine the effect of this structural change on the energetics of K^+^ ion permeation we used the same umbrella sampling protocol to obtain the PMF profile for a K^+^ ion permeation through the nhTMEM16 L302A mutant in a 3:1 POPE/POPG membrane system. The groove of the mutated scramblase remains in its original conformation during the umbrella sampling simulation (Supplementary Figure S9) and there is no lipid in the groove.

Figure 5 shows the PMF profile of K^+^ permeation in the L302A system with the ranges of “critical interactions” marked by the double arrows. The free energy barrier of the K^+^ ion permeation for this construct is ∼8.0 kcal/mol, similar to the value calculated for System 3 in which a POPC lipid resided in the open groove of the wild type nhTMEM16. The maximum is at Z = 3.5 Å, where the structure shows the pore to be narrowest, and the translocated K^+^ ion encounters the hydrophobic residue L336, which dynamically occludes the pathway (Supplementary Figure S10). The EC gate in the simulation of this construct is mostly closed as E313 interacts with R432 (Supplementary Figure S11). Nonetheless, E313 still acts as an attracting site for the K^+^ ion, as indicated by the location of the minimum in the PMF curve near the range of the critical interaction with (Figure 5). We note that a POPG at the EC gate interacts with R432 and the K^+^ ion (Supplementary Figure S12), as observed also in the unbiased simulation of wild type.

**Figure 5:**
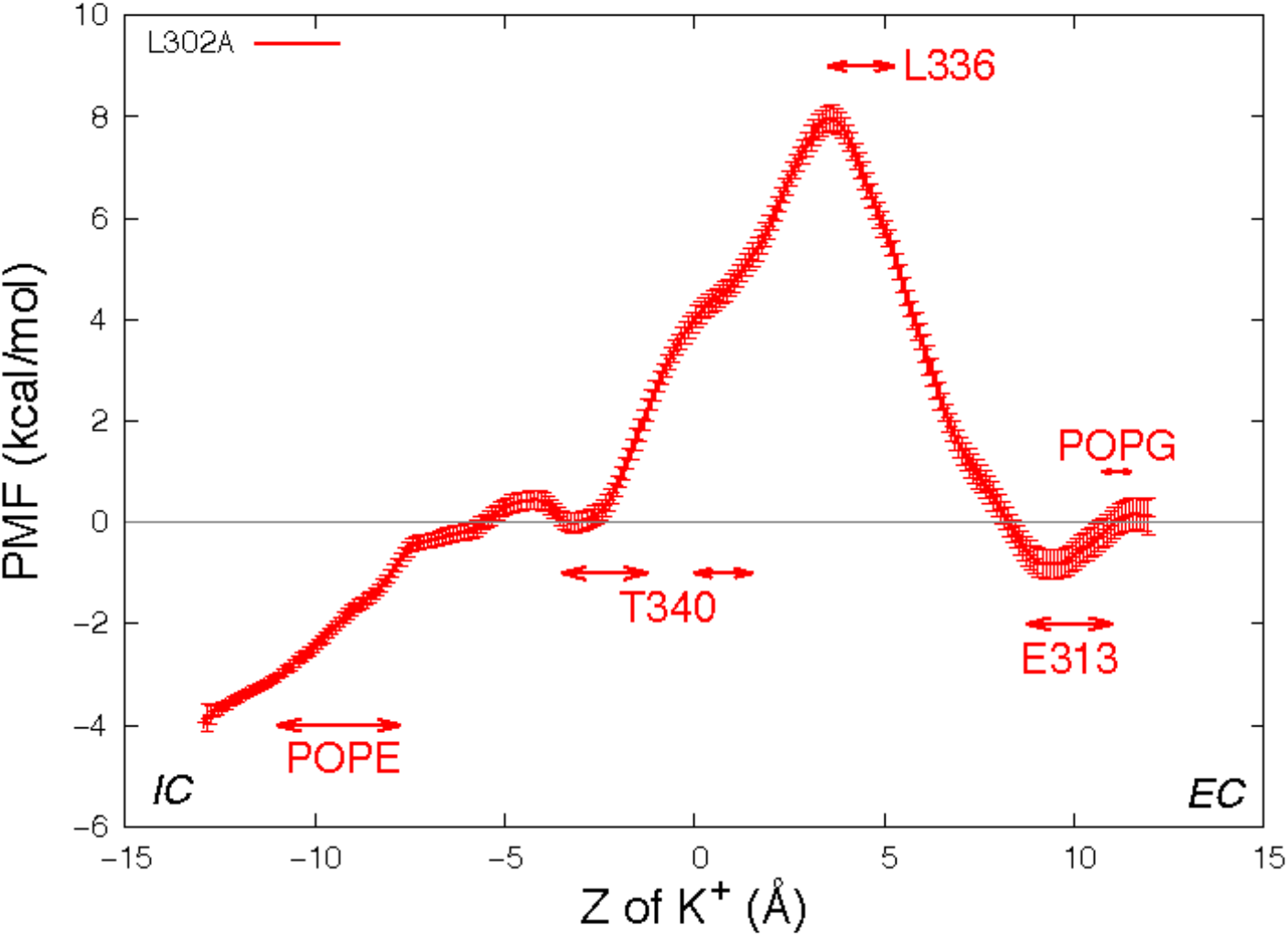
Potential of mean force profile of a K^+^ ion along the permeation pathway in the groove of one protomer of the nhTMEM16-L302A mutant (see Methods for the description of the error bar calculations). Z = 0 is set to be the Z position of the T381 Cα atom of this subunit. As in Fig 3, the “critical interaction” ranges are marked by horizontal double arrows.

### Electrostatic properties of the nhTMEM16 groove play mechanistic role in regulating energetics of the K^+^ ion permeation process

The space sampled by the K^+^ ion as it progresses along the permeation path is summarized in Figure 6 where the purple spheres indicate positions visited by the potassium ion in all the frames from the equilibrated umbrella sampling simulations. For System 1, the groove volume sampled by the potassium ion does not enclose the whole POPG head group, as the ion passes the in-groove lipid head group “on the side” (see details in the “zoomed-in” insert), without being blocked by it. However, for System 3, the head group of POPC is immersed in the volume of sampling ions, which supports the inference from the corresponding PMF profile that this lipid hinders the potassium ion permeation. Overall, the space sampling is very similar in all wild type nhTMEM16 constructs with open-groove conformations, but different at the narrow “neck” of the groove in the L302A where the side chains of Y439 and L336 form a constriction (see Supplementary Figure S10).

**Figure 6:**
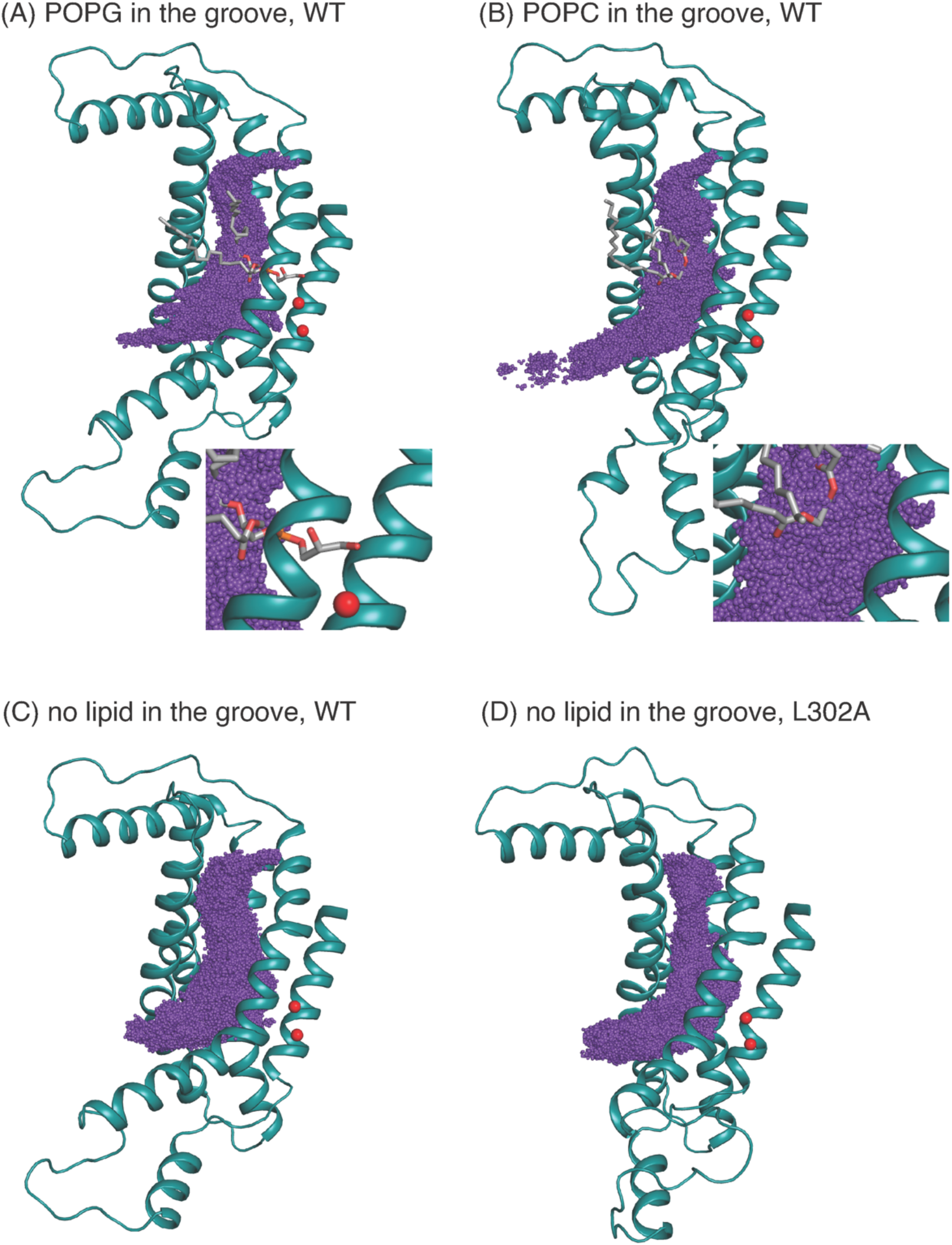
K^+^ positions from all equilibrated frames of the umbrella sampling simulation indicated by purple spheres in a structural representation of the groove in which only TM3–TM7 are shown for clarity, rendered in teal cartoon. The in-groove lipid is rendered in licorice, and Ca^2+^ ions are shown as red spheres. Panel (**A**) is for System 1; Panel (**B**) is for System 3, Panel (**C**) is for System 2, and Panel (**D**) is for the nhTMEM16 L302A mutant. System 4 is similar to System 2 and therefore not shown. The insets in panels A and B show an enlarged view of the sampling near either the POPG (panel A) or POPC (panel B).

The “average trajectory” paths for the K^+^ ion translocation in the grooves of the various nhTMEM16 constructs (shown in Figure 7A, B) were obtained by averaging the XY coordinates of the positions sampled by the K^+^ ion (purple spheres in Figure 6) at a particular Z. These trajectories are seen to be closely aligned with the groove axes obtained with the HOLE software ^24^ (http://www.holeprogram.org/) which considers only geometric measures of the molecular components (i.e., atom radii). The paths of the translocated ion shown in Figure 7A-B are seen to follow the groove axis enabling a comparison of the steric environment in the WT and L302A systems using HOLE ^24^ (see Figure 7C, D) and of the electrostatic potential (EP) along the “averaged trajectories” (Fig. 8). The values of EP along the trajectories in the four WT constructs and the L302A mutant of nhTMEM16 (Figure 8) were calculated by solving the linearized Poisson-Boltzmann equation ^25^ in CHARMM ^26^ (see Methods for details) for only one static frame of the protein (the starting configuration). The results showing that the EP in the nhTMEM16 groove region can be positive or negative in various regions of the different constructs are consistent with the experimental findings that their ion channel activity is not selective for cations or anions ^27^. The generally slightly negative value of the electrostatic potential in Systems 2 and 4, is enhanced by the presence of the POPG lipid in the groove (System 1), consonant with the relatively low free energy barriers calculated for the potassium ion permeation. In contrast, the presence of a POPC lipid in the groove of the wild type nhTMEM16 (System 3), and the narrowing of the groove in the L302A mutant, result in an overall positive EP, which is further consistent with our finding relatively high free energy barriers for the potassium ion translocation in these systems (∼7-8 kcal/mol).

**Figure 7:**
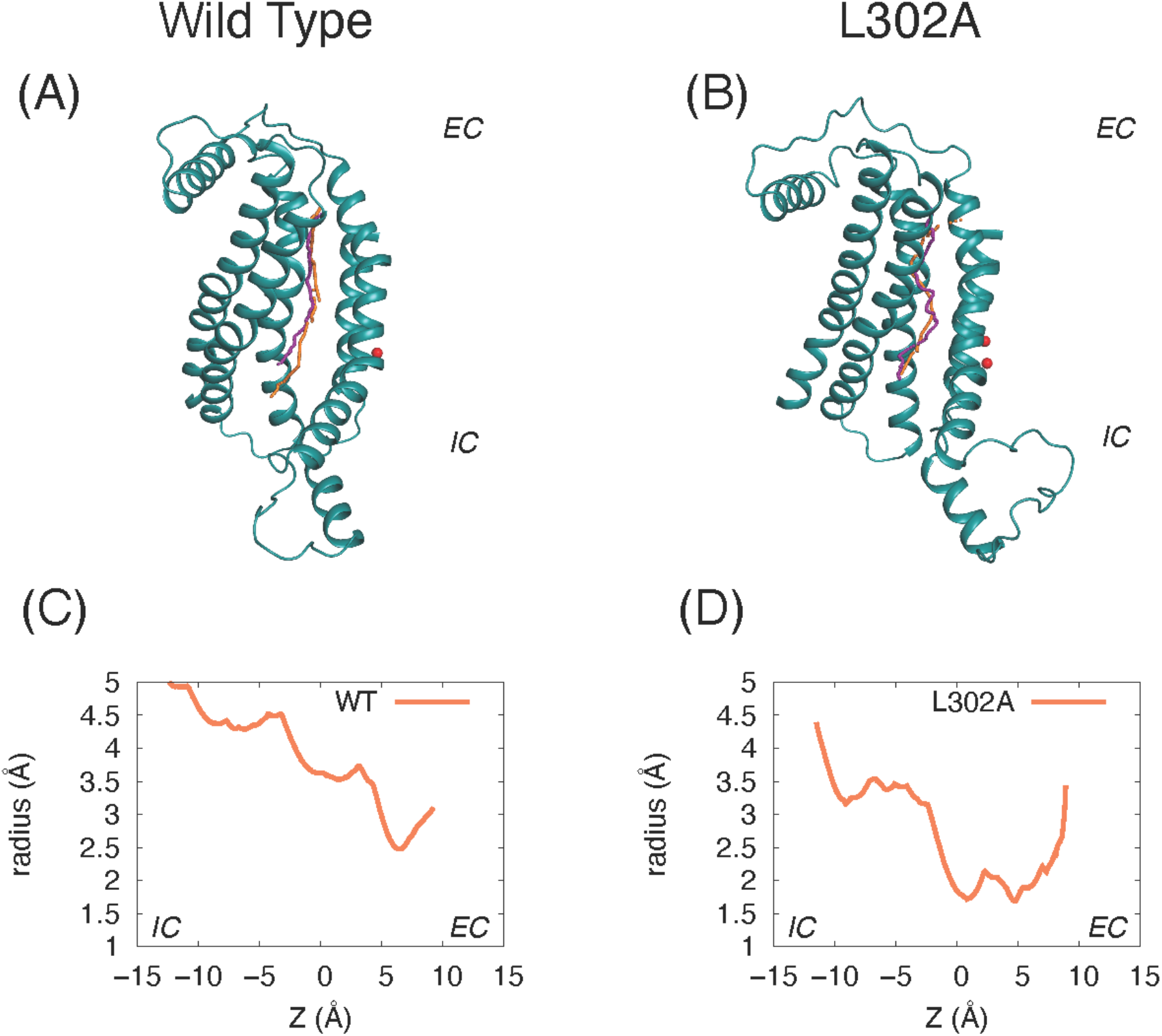
Comparison of the groove axis obtained from the Hole calculations to the “average K^+^ trajectory” obtained by averaging the XY positions of K^+^ at each Z value in the umbrella sampling calculations. Panels (**A**) and (**C**) are from System 2 calculations of the wild type nhTMEM16 in a 3:1 POPE/POPG membrane. Panels (**B**) and (**D**) are from parallel simulations of the nhTMEM16-L302A mutant. The groove axis calculated with the HOLE software is in orange, and the K^+^ trajectory in purple. For simplicity, only the protein region composed of the TM3–TM7 helices is shown (rendered in teal cartoon), and Ca^2+^ ions are shown as red spheres. Panels (C) and (D) show the radius of the pore along the pathway calculated with the HOLE software as described above.

**Figure 8:**
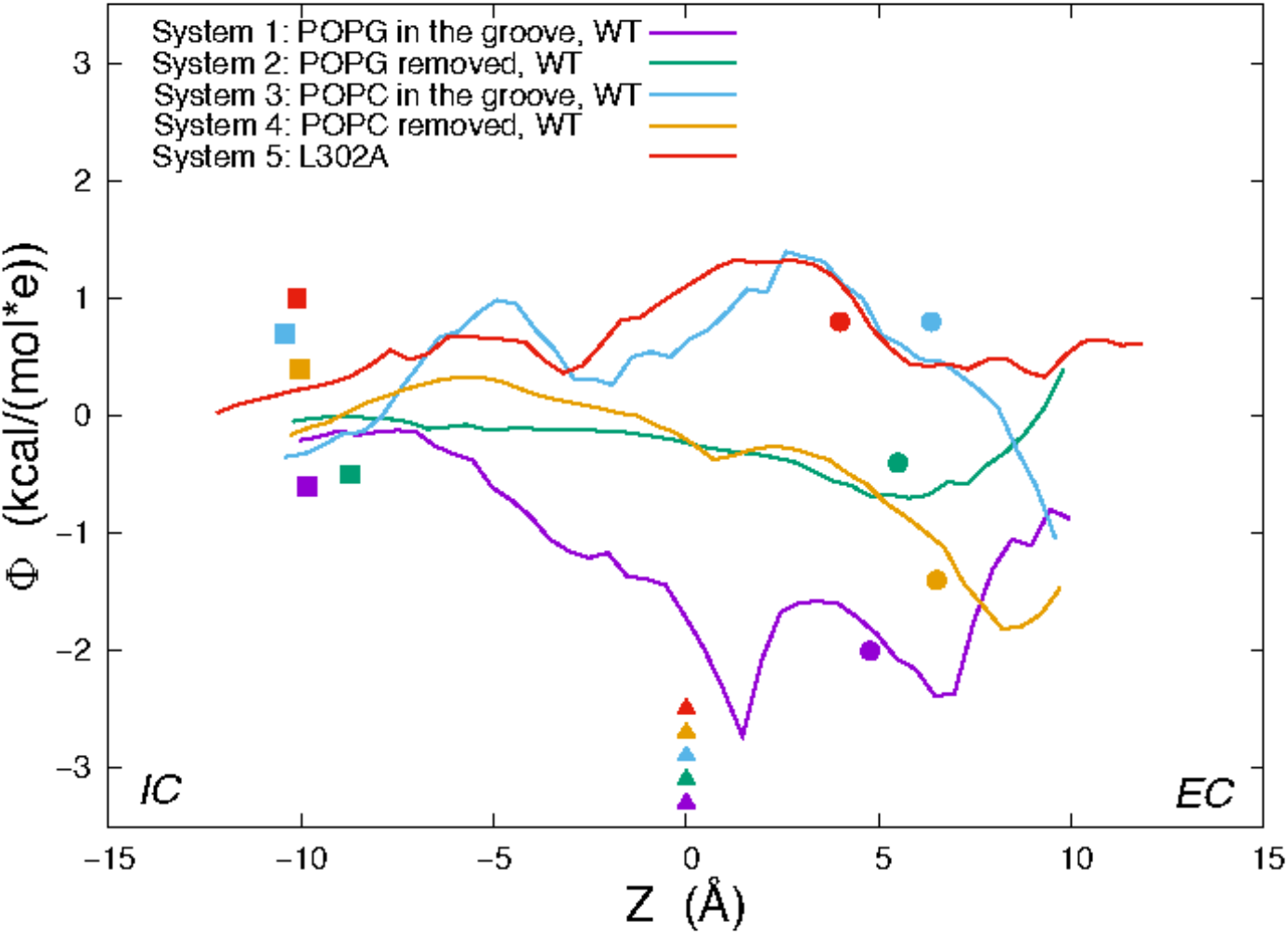
Electrostatic potential (EP) values calculated along the average trajectory of K^+^ from umbrella sampling (see text and Figure 6) for the wild type nhTMEM16 Systems 1– 4 and the nhTMEM16-L302A mutant. For orientation along the trajectories, the colored symbols indicate the Z positions of the following landmarks: C_α_ atom of T381 (triangles); C_α_ atom of Y439 (circles); the highest placed Ca^2+^ ion (squares).

## Discussion

The permeation events observed from our conventional MD simulations, with no voltage applied, show that potassium ions can pass through the hydrophilic groove of the nhTMEM16 scramblase, consistent with the reports that it can function as a non-selective ion channel ^27^. But the translocation efficiency is dependent on the presence of lipids in the groove, as suggested by the potential of mean force calculations for the permeation paths under various conditions. When the protein is in the Ca^2+^-activated state, the presence of lipids that are being scrambled or are diffusing in and out of the groove as observed in simulations, can reduce potassium ion permeation to various extents. To interpret the mechanistic reasons for the calculated free energy profile for the translocation, we identified the effects of specific residues involved in K^+^ permeation along the ion path. These include multiple polar residues, such as E313 (in the EC gate region ^19^), T333, T381, and S382 (see Fig. 1B), as well as the in-groove POPG lipid – all of which generate favorable conditions for the ions to pass through the open groove conformation of nhTMEM16. In contrast, the presence of electrostatically neutral POPC lipid in the groove creates a free energy barrier to ion movement, which is ∼5 kcal/mol higher than the in-groove POPG.

To consider the role of the groove 3D topography we carried out parallel calculations for the nhTMEM16 mutant L302A, in which groove occlusions prevent lipid scrambling even in the presence of Ca^2+ 14^. In the L302A mutant, the free energy barrier for the K^+^ translocation is found to be 6 kcal/mol higher than in the WT construct, and a significantly lower K^+^ permeation under the same condition (Fig. 4). While we found that the EP calculated in the grooves of the four WT systems simulated contains both positive and negative patches, indicating that the ion channel is not charge selective, the profile of the electrostatic potential in the permeation path of the L302A mutant is different and suggests that anion permeation may be favored.

In our unbiased 10 µs long MD simulation of the nhTMEM16 wild type in 3:1 POPE/POPG membrane system, we did not observe any instance of chloride passing through the groove. Notably, the number of Cl^−^ ions in the simulation box is much smaller than the number of potassium ions (18 Cl^−^ versus 223 K^+^), because the membrane is negatively charged, reducing the probability of a successful Cl^−^/protein encounter. Indeed, our unbiased 15 µs long MD simulation of nhTMEM16 in POPC membrane system ^14^, where the number of Cl^−^ is similar to that of K+ (124 Cl− and 130 K^+^), we did observe spontaneous Cl^−^ ion penetration into the groove. However, coming from the IC end and moving up only to the middle of the groove, the ion did not complete the translocation in this run, precluding the initiation of a comparative umbrella sampling simulations for Cl^−^ similar to those for K^+^. In future simulations of much longer times this permeation is expected to be completed and thereby enable full comparative evaluations.

## Methods

We have previously described in detail the unbiased MD simulation protocols used here for the nhTMEM16 wild type and the L302A mutant 14,19. Here, the 3:1 POPE/POPG membrane containing the wild type nhTMEM16 was constructed of 851 lipids and a solution box containing 86,011 waters and 150 mM K+Cl-, resulting in a total of 388,447 atoms. The POPC membrane containing the wild type scramblase had 679 lipids in a solution box of 90,189 water molecules and 150 mM K+Cl-, for a total of 385,207 atoms. The 3:1 POPE/POPG membrane containing the L302A mutant had 1505 lipids and 146,737 waters in addition to the 150 mM K+Cl-, resulting in a total of 652,878 atoms. All the simulations implemented the CHARMM36 force-field parameters 28. Umbrella sampling simulations (see below) were performed using NAMD software version 2.13 (28), with a 2 fs integration time-step, under the NPT ensemble (at T = 310 K).

### Umbrella Sampling

#### System preparation

Before umbrella sampling, Systems 2 and 4 (WT nhTMEM16 with no lipids in the groove) were equilibrated for 11.8 ns and 6.2 ns after the in-groove lipid was removed from Systems 1 and 3 respectively. Because ion permeation is generally much faster than lipid translocation (104 lipids per second 3), 1D umbrella sampling calculations of ion permeation were carried out for all systems with and without a lipid in the groove. The head group of the POPC in the groove of System 3 is at about the same height (Z position along the membrane axis) as the head group of the in-groove POPG of System 1, shown in Supplementary Figure S3. This was achieved by identifying a frame from our earlier unbiased MD simulations of nhTMEM16 in the POPC membrane system, in which there was a POPC lipid at about the same height in the groove, and the EC gate was broken. This POPC lipid was not scrambled either but diffused towards the middle of the groove from IC, and then moved back to IC. It is worth mentioning that there are always lipids in the IC region of the groove; here a lipid is called “in-groove” if it is situated in the middle of the groove, near or above V447, which is positioned in the middle of TM6 and faces the groove (see Supplementary Figure S8); when we removed the “in-groove” lipid, some lipids in the IC region can move up along the groove, but on the relatively short timescales of the umbrella sampling simulations (2.4 ns), they don’t move high up the groove to be considered as the “in-groove” lipids in the umbrella sampling simulations.

Potential of Mean Force calculations: The Z position of the potassium ion relative to the Z position of the Cα atom of T381 was chosen as the reaction coordinate for the calculation. The residue T381 in TM5 was chosen as it is positioned in the middle of the transmembrane region of the protein (Supplementary Figure S8), and its dynamic fluctuations are relatively small during simulations. We defined 84 umbrella sampling windows covering Z = -10.25 Å to Z = 10.5 Å range (Z = 0 Å representing the position of the Cα atom of T381), with a spacing of 0.25 Å between the centers of neighboring constraints. The bias is a harmonic potential function with a force constant equal to 10 kcal/mol/Å2. Additional windows were added with force constant of 20 kcal/mol/Å2 to improve the sampling for Z = 3.6 Å window in System 1 (see Supplementary Figures 13-14 for overlaps of umbrella sampling histograms). For each window, we performed 5 independent runs of 2.4 ns, including 0.6 ns equilibration that was discarded for further analysis. The initial structural configurations for umbrella sampling windows are taken from the unbiased simulation frames where we observed a K+ ion in the groove. The potential of mean force was calculated using Weighted Histogram Analysis Method (WHAM) 29,30, and Monte Carlo bootstrap error analysis was carried out within the WHAM program by setting up a correlation time of 20 ps.

For the mutant nhTMEM16 L302A system, the initial structural configuration is from a frame at the end of a 2 µs unbiased MD simulation of the system. To obtain starting configurations for all the umbrella sampling windows for this system, a steered MD simulation was performed with force constant of 5 kcal/mol/Å2 and movement velocity of 10 Å/ns. The setup of the umbrella sampling for the L302A system was the same as the one used for the wild type nhTMEM16, except that we used 96 umbrella sampling windows covering Z = -12.25 Å to Z = 11.5 Å range.

### Electrostatic Potential Calculations

The electrostatic potential was calculated by solving the linearized Poisson-Boltzmann equation ^25^ in CHARMM ^26^. The protein was assigned a dielectric constant of E_*P*_ = 2. Its TM region was embedded in a membrane slab, which includes a 26-Å-thick hydrophobic core of the bilayer (with E_*core*_ = 2 dielectric constant), surrounded by two 8-Å-thick regions, representing the lipid headgroups (E_*head*_ = 30). The membrane slab contained a cylindrical hole (with the dielectric of E_*W*_ = 80) with a radius of 18 Å centered around the groove of the protomer. The system was surrounded by solvent (E_*W*_ = 80) containing 150 mM of monovalent mobile ions. The cylindrical hole was accessible to the ions, but ions were not permitted into the membrane region.

## ACKNOWLEDGMENTS

We gratefully acknowledge stimulating discussions with Prof. Alessio Accardi and member of his lab. This work was supported by National Institutes of Health (NIH) Grant R01GM106717. GK and HW acknowledged support from the *1923 Fund*. The following computational resources are gratefully acknowledged: the Anton 2 supercomputer at the Pittsburgh Supercomputing Center (PSC) provided through Grant R01GM116961 from the National Institutes of Health (The Anton 2 machine at PSC was generously made available by D.E. Shaw Research); resources of the Oak Ridge Leadership Computing Facility (INCITE allocation BIP109) at the Oak Ridge National Laboratory, which is supported by the Office of Science of the U.S. Department of Energy under contract no. DE-AC05-00OR22725; and the computational resources of the David A. Cofrin Center for Biomedical Information in the Institute for Computational Biomedicine at Weill Cornell Medical College.

