## Supplementary Material for "The permeation of potassium ions through the lipid scrambling path of the membrane protein nhTMEM16"

### Supplementary FIGURES

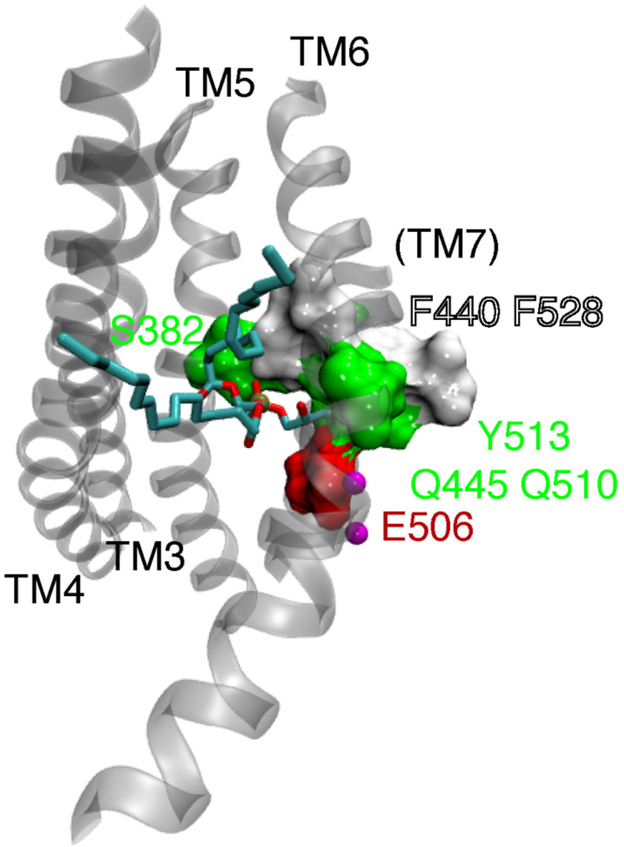


**Figure S1**: A POPG lipid in the groove of the wild type nhTMEM16 in 3:1 POPE/POPG membrane system. Within 3Å of the POPG head group are S382 (TM5), F440 (TM6), Q445 (TM6), E506 (TM7), Q510 (TM7), Y513(TM7) and F528 (TM8), shown as surface and colored according to residue type: hydrophobic as white, polar as green, negatively charged as red. POPG is shown as licorice. TM3-TM7 of the protein are shown as transparent cartoon, and $Ca^{2+}$ ions are shown as purple spheres.

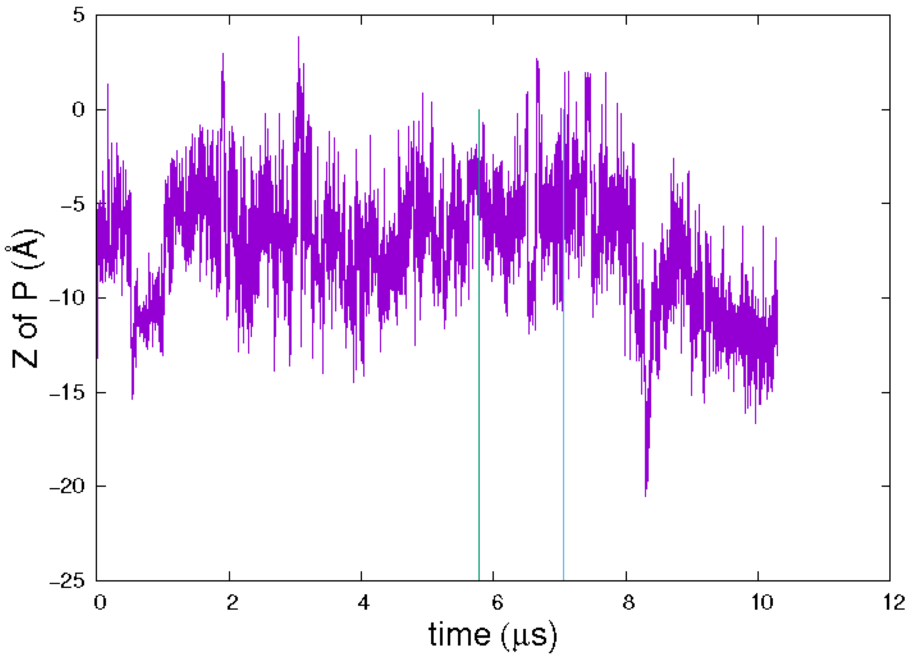


**Figure S2**: Z position of the phosphorous atom (P) of the groove-bound POPG (relative to the Z position of the $C_\alpha$ atom of residue T381), during the unbiased 10 μs MD simulation of the wild type nhTMEM16 in 3:1 POPE/POPG membrane system. Vertical lines depict when the two $K^+$ permeation events occur.

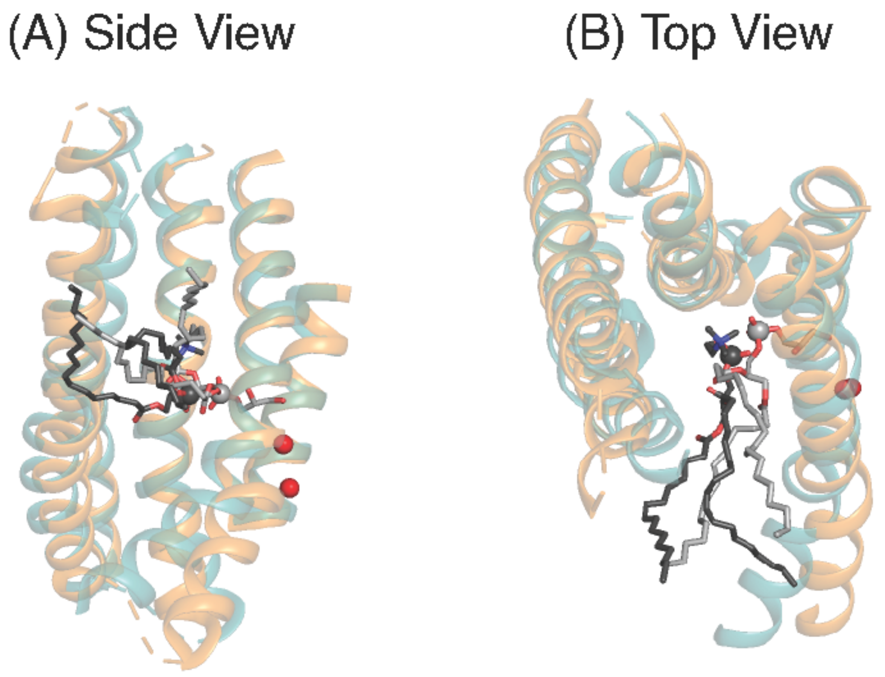


**Figure S3**: Alignment of the wild type nhTMEM16 in two simulation systems: (1) 3:1 POPE/POPG membrane system with a POPG in the groove and (3) POPC membrane system with a POPC in the groove. The POPG lipid is shown as light grey licorice, the POPC lipid is shown as dark grey licorice, and phosphorus atoms are shown as spheres. TM3{TM7 of one subunit are shown as teal cartoon for system (1) and as orange cartoon for system (3). $Ca^{2+}$ ions of system (1) are shown as red spheres.

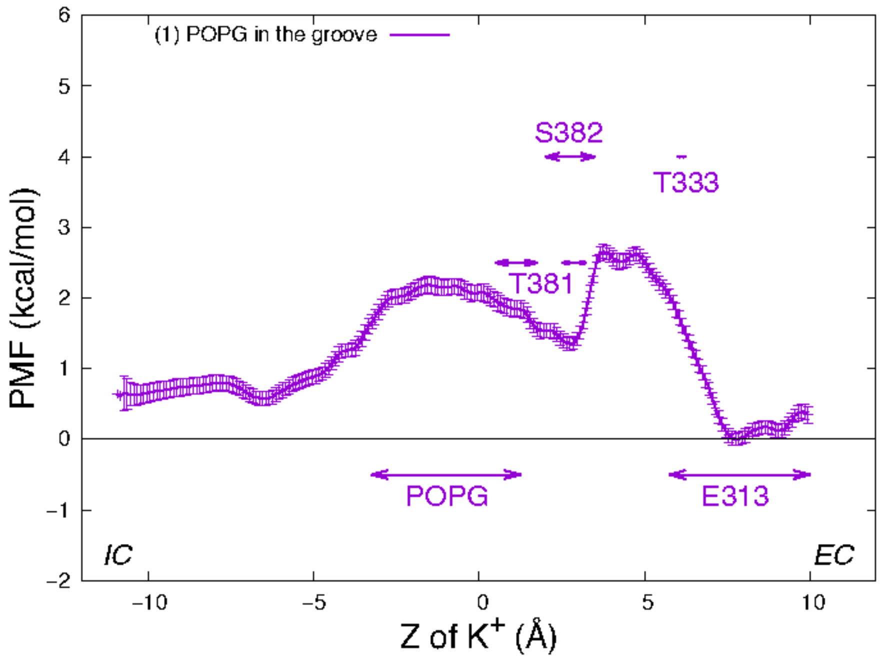


**Figure S4**: Potential of mean force of a $K^+$ ion along the permeation pathway in system (1), wild type nhTMEM16 in 3:1 POPE/POPG membrane system with a POPG in the groove. PMF is depicted with error bars. Z = 0 is set to be the Z position of T381 $C_\alpha$ atom of this subunit. The interaction ranges of the K+ ion with \critical" residues are marked by horizontal double-arrowed lines.

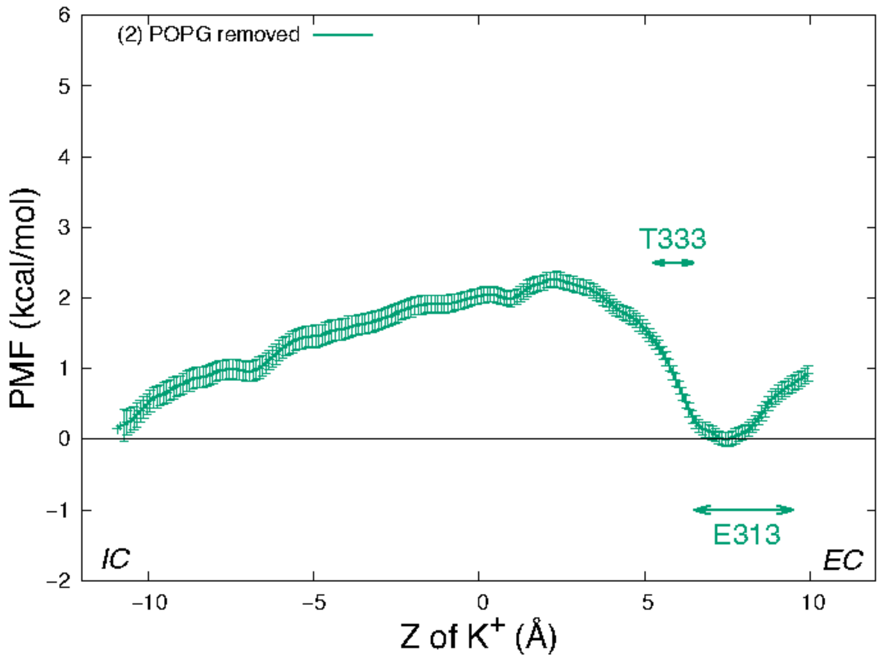


**Figure S5**: Potential of mean force of a K+ ion along the permeation pathway in system (2), wild type nhTMEM16 in 3:1 POPE/POPG membrane system, without a lipid in the groove. PMF is depicted with error bars. Z = 0 is set to be the Z position of T381 $C_\alpha$ atom of this subunit. The interaction ranges of the K+ ion with \critical" residues are marked by horizontal double-arrowed lines.

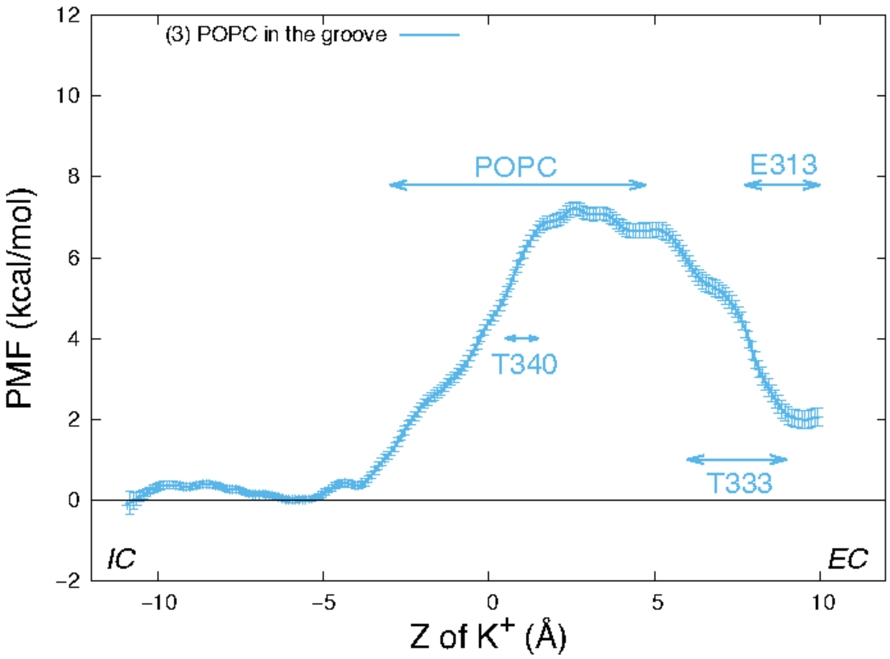


**Figure S6**: Potential of mean force of a K+ ion along the permeation pathway in system (3), wild type nhTMEM16 in POPC membrane system with a POPC in the groove. PMF is depicted with error bars. Z = 0 is set to be the Z position of T381 $C_\alpha$ atom of this subunit. The interaction ranges of the K+ ion with \critical" residues are marked by horizontal double-arrowed lines.

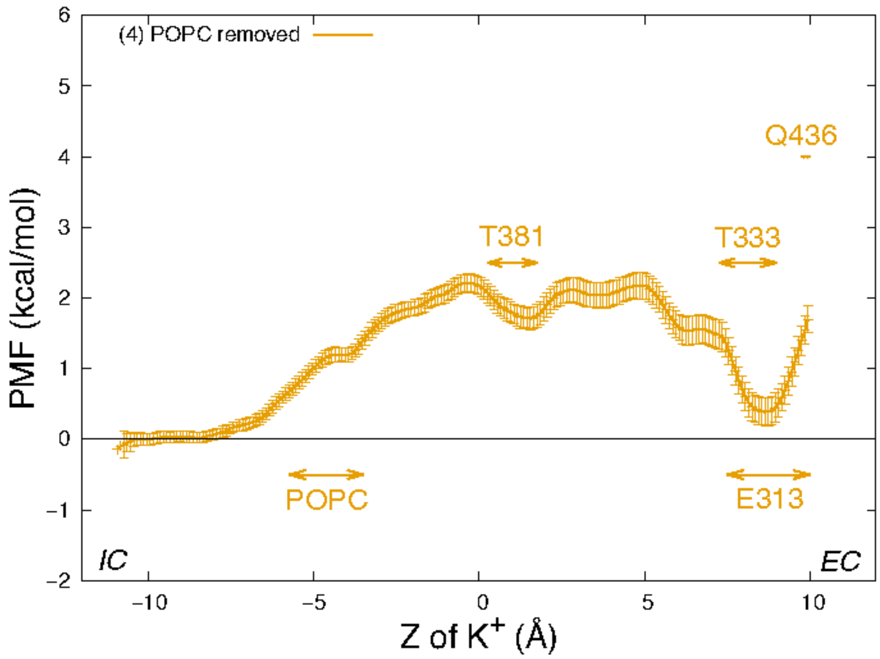


**Figure S7:** Potential of mean force of a K+ ion along the permeation pathway in system (4), wild type nhTMEM16 in POPC membrane system, without a lipid in the groove. PMF is depicted with error bars. Z = 0 is set to be the Z position of T381 $C_\alpha$ atom of this subunit. The interaction ranges of the K+ ion with \critical" residues are marked by horizontal double-arrowed lines.

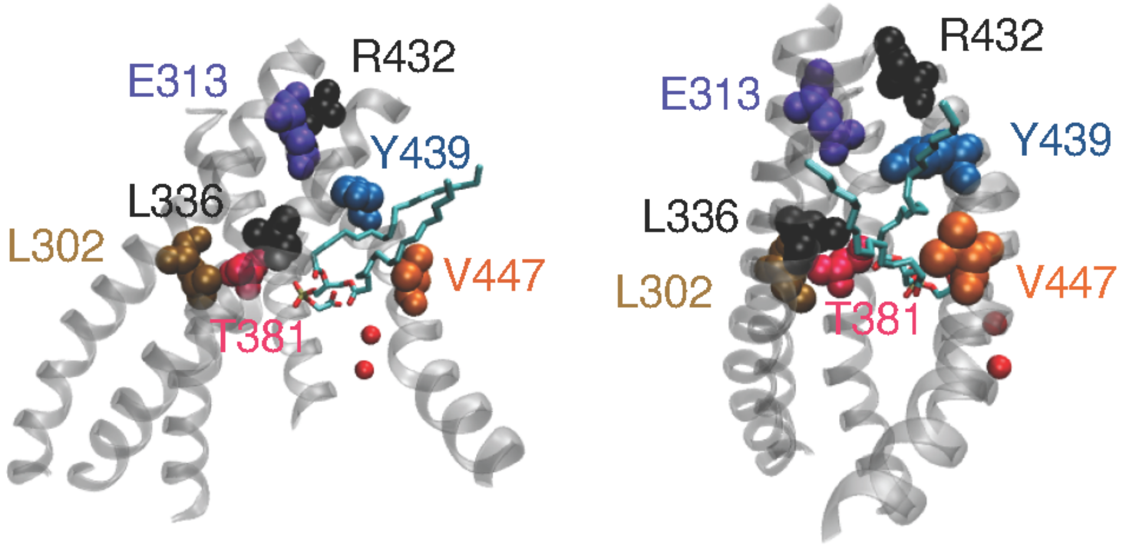


**Figure S8:** Two perspective views of the groove of the wild type nhTMEM16. Residues shown are: L302, E313 of TM3; L336 of TM4; T381 of TM5; R432, Y439, and V447 of TM6.

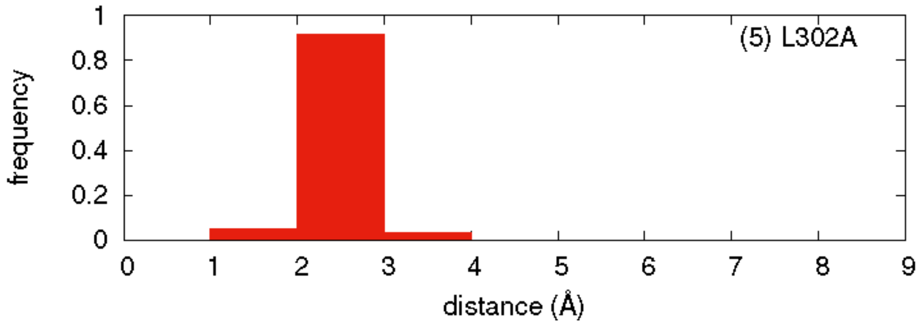


**Figure S9:** Histogram of the distance between V337 and V447 of the nhTMEM16 L302A mutant. Minimal distance between any atom of V337 and any atom of V447 was calculated from the equilibrated frames from all windows in the umbrella sampling simulations.

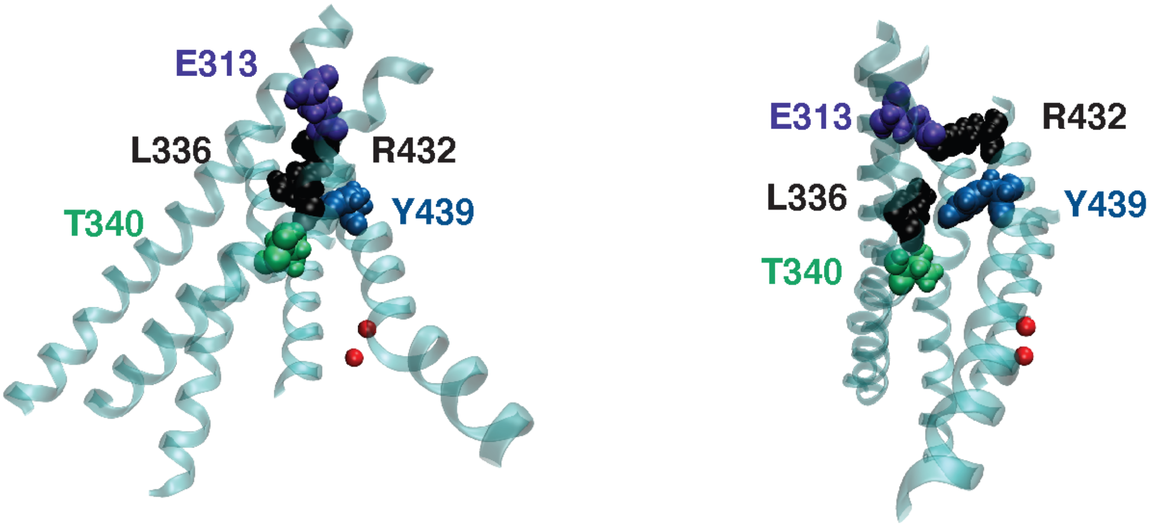


**Figure S10:** Two perspective views of the groove of the nhTMEM16 L302A mutant. Residues shown are: E313 of TM3; L336 and T340 of TM4; R432 and Y439 of TM6.

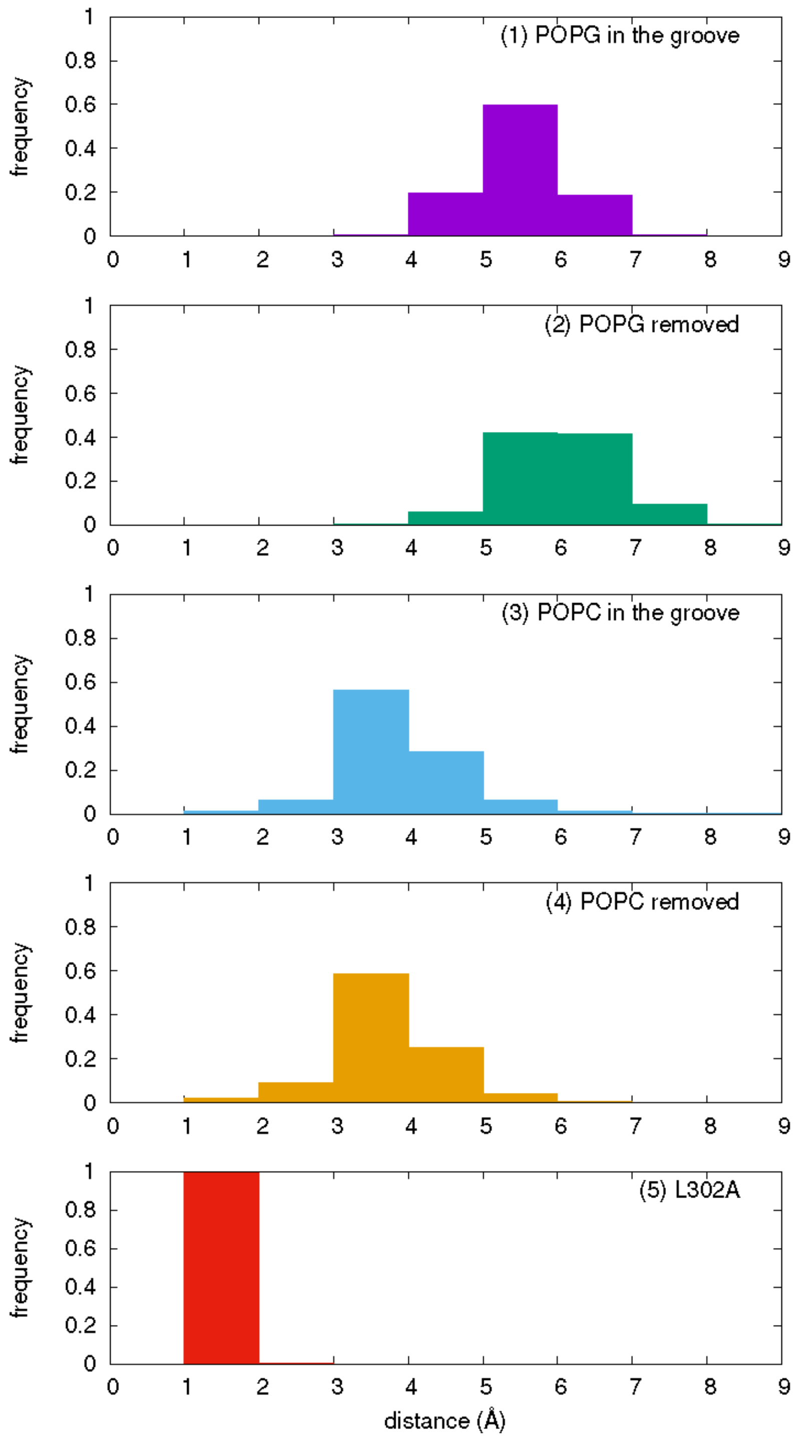


**Figure S11:** Histogram of the distance between E313 and R432 of nhTMEM16 in different systems investigated in the main text. Minimal distance between any atom of E313 and any atom of R432 was calculated from the equilibrated frames from all windows in the umbrella sampling simulations.

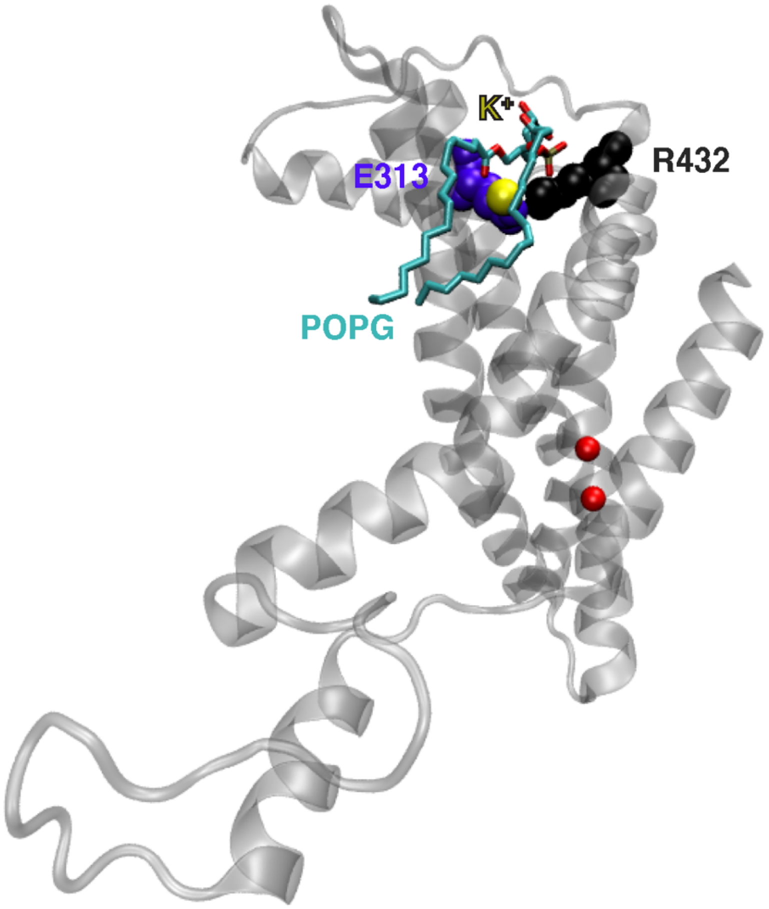


**Figure S12:** Four residues interacting at the EC of nhTMEM16 L302A mutant. E313 and R432 are represented as spheres. POPG 47 is represented as licorice, and $K^+$ ion is represented as yellow sphere.

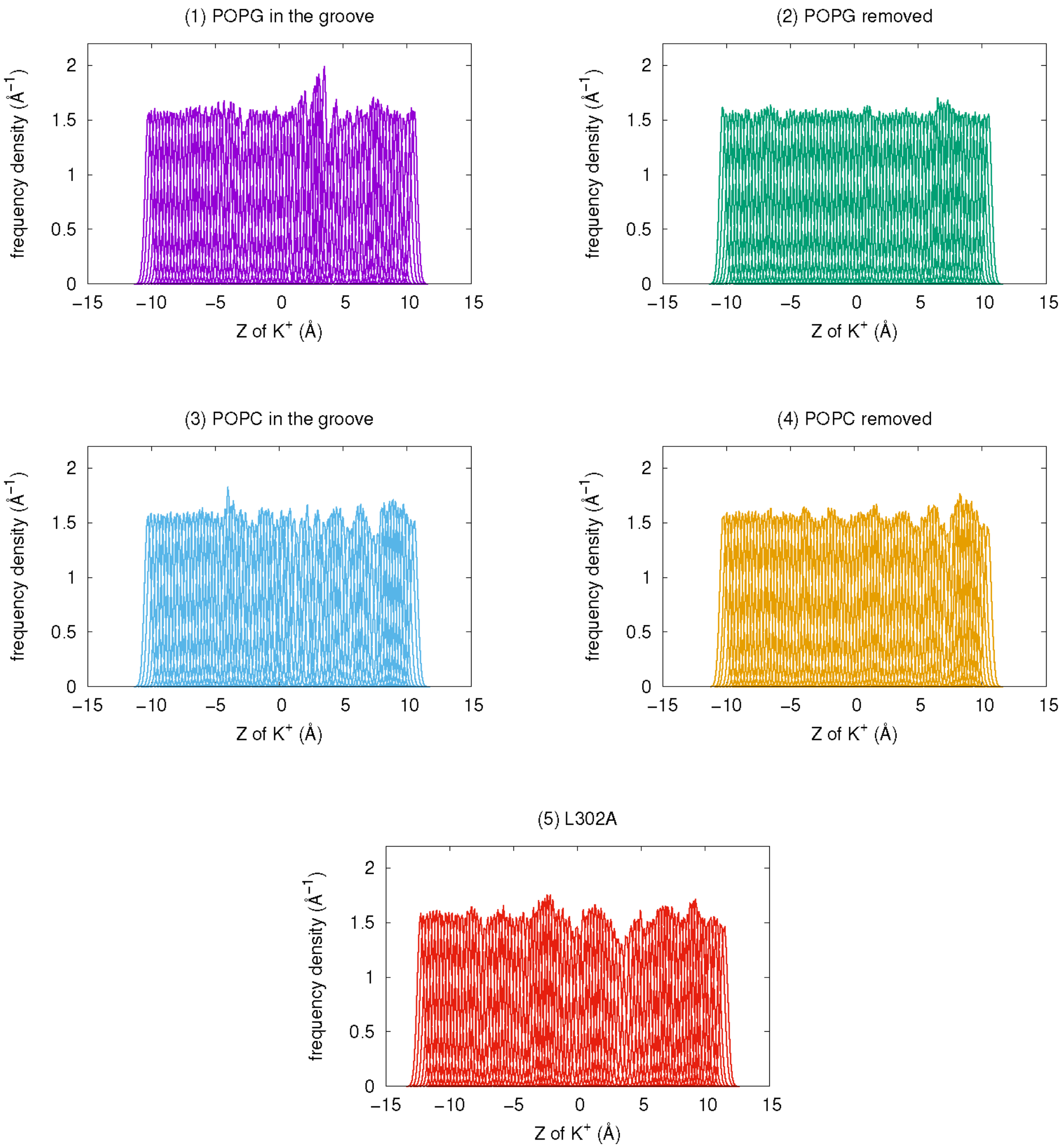


**Figure S13:** Overlap of umbrella sampling histograms for the five simulated systems.

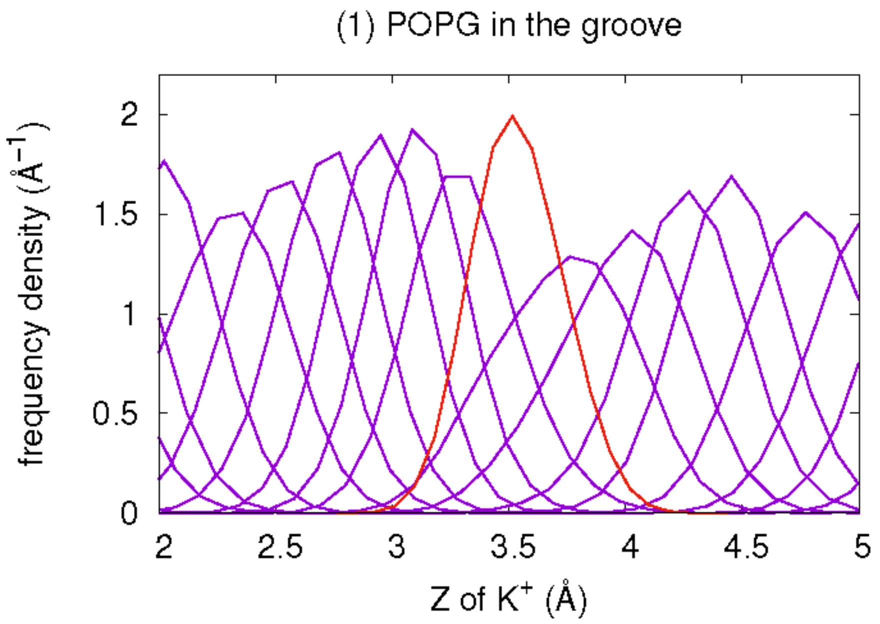


**Figure S14:** Overlap of umbrella sampling histograms for System 1, zoomed in the 2Å to 5Å interval of Z coordinate. An additional window was added at Z = 3.6Å to improve the sampling (depicted as red).
